# Structural insights into HIV-1 polyanion-dependent capsid lattice formation revealed by single particle cryo-EM

**DOI:** 10.1101/2022.12.02.518872

**Authors:** Carolyn M. Highland, Aaron Tan, Clifton L. Ricaña, John A.G. Briggs, Robert A. Dick

## Abstract

The HIV-1 capsid houses the viral genome and interacts extensively with host cell proteins throughout the viral life cycle. It is composed of capsid protein (CA), which assembles into a conical fullerene lattice composed of roughly 200 CA hexamers and 12 CA pentamers. Previous structural analyses of individual CA hexamers and pentamers have provided valuable insight into capsid structure and function, but high-resolution information about these assemblies in the broader context of the capsid lattice is lacking. In this study, we combined cryo-electron tomography and single particle analysis cryo-electron microscopy to determine high-resolution structures of continuous regions of the capsid lattice containing both hexamers and pentamers. We also developed a new method of *in vitro* lattice assembly that enabled us to directly study the lattice under a wider range of conditions than has previously been possible. Using this approach, we identified a critical role for inositol hexakisphosphate (IP6) in pentamer formation and determined the structure of the CA lattice bound to the capsid-targeting antiretroviral drug GS-6207 (Lenacapvir). Our work reveals new structural details of the mature HIV-1 CA lattice and establishes the combination of lattice templating and single particle analysis as a robust strategy for studying retroviral capsid structure and capsid interactions with host proteins and antiviral compounds.

**Significance statement:** The mature HIV-1 capsid is composed of the capsid (CA) protein arranged in a conical lattice of hexamers and pentamers. Numerous structures of individual CA hexamers and pentamers alone have been published, but the molecular details of these assemblies in a more global, lattice-wide context are lacking. Here, we present high-resolution cryo-electron microscopy structures of continuous regions of the capsid lattice containing both hexamers and pentamers. We also describe key differences in the assembly and structures of these oligomers that have important implications for understanding retroviral maturation and for ongoing efforts to pharmacologically target the HIV-1 capsid.

## INTRODUCTION

After budding from a cell, immature HIV-1 undergoes maturation to become infectious. In this viral protease-mediated process, the incomplete spherical immature protein lattice of ∼400 Gag hexamers is dismantled and replaced by a new, morphologically distinct mature lattice of capsid (CA) molecules proteolytically released from Gag (Supplemental Figure 1A) (reviewed in (1, 2). CA assembles into a conical lattice formed by roughly 200 CA hexamers and exactly twelve CA pentamers, the latter of which facilitate lattice curvature and closure around the viral genome (3–5). Monomeric CA consists of two folded domains connected by a flexible linker (6–8): the N-terminal domain (NTD) faces the outer surface of the capsid, where it stabilizes the hexamer, while the C-terminal domain (CTD) forms dimeric interactions that link hexamers together (4, 9, 10). Beyond its structural and genome-protective roles, the capsid also interfaces extensively with host cell proteins, many of which have been co-opted by the virus to ensure its proper trafficking and uncoating within the cell. As such, CA has become the focus of an emerging class of capsid-targeting antiretroviral compounds that interfere with capsid-host factor interactions and compromise capsid structural integrity (reviewed in (11–13).

**FIGURE 1.**
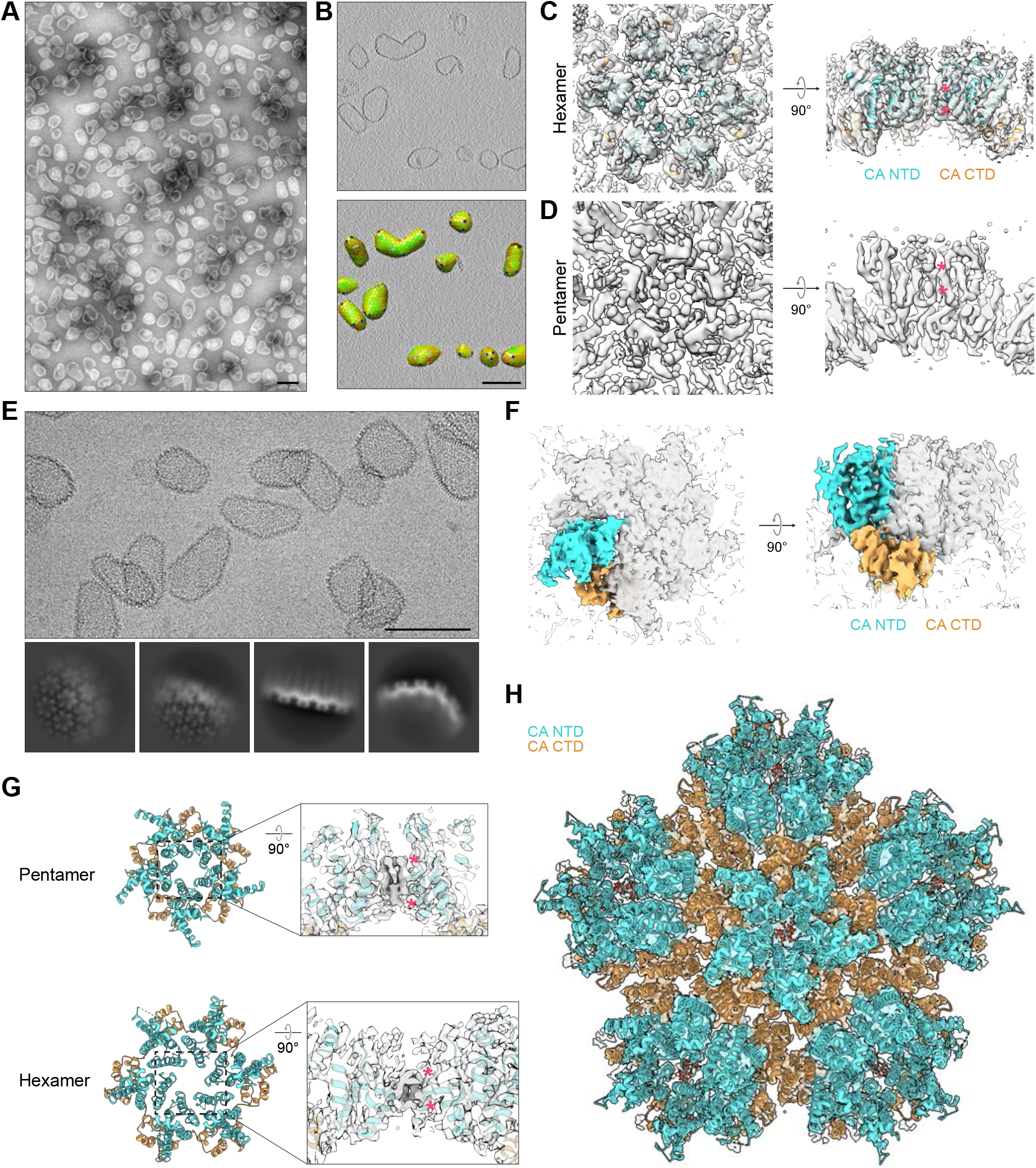
Cryo-electron tomography and single particle cryo-electron microscopy of HIV-1 CA capsid-like particles. **A**. Negative stain TEM micrograph of capsid-like particles (CLPs) prepared *in vitro* from purified CA and IP6. **B**. Central ‘slice’ through CLPs in a tomogram (top) with corresponding lattice maps determined by subtomogram averaging (bottom). Hexamers in the lattice map are colored from green to red according to cross-correlation (CC) with the average hexamer structure, with red indicating lower CC and green indicating higher CC. Pentamers are colored blue. **C**. Cryo-ET reconstruction of the CLP hexamer with crystal structure 4XFX (Gres et al., 2015) docked according to best cross-correlation with the cryo-EM density. Densities corresponding to IP6 are marked with magenta asterisks. **D**. Cryo-ET reconstruction of the CLP pentamer. Densities corresponding to IP6 are marked with asterisks. **E**. Cryo-EM micrograph of CLPs (left) with select 2D class averages (right). **F**. Cryo-EM (SPA) map of CLP central pentamer. Density corresponding to a single CA monomer is highlighted with color. **G**. Right: densities corresponding to IP6 (marked with magenta asterisks) in pentamer (top) and hexamer (bottom) cryo-EM maps. Left: atomic model for CLP pentamer (top) and hexamer (bottom). **H**. Atomic model for ‘global’ pentamer structure (pentamer surrounded by its five nearest hexamer neighbors) with corresponding cryo-EM map. *myo*-IP6 molecules are docked according to best cross-correlation with the cryo-EM density. In all maps and models shown, the CA NTD is shown in cyan and the CA CTD is shown in orange. Side views in C, D, and G show central pore-cross sections. Scale bars, 100 nm.

Most published high-resolution structural information about the mature capsid has come from x-ray crystallography and NMR analyses of individual CA hexamers, pentamers, and monomers. Although these studies have certainly provided invaluable insights into capsid structure (8, 14–18) and its interactions with small molecules (reviewed in (12) and host proteins (6, 19–21), we currently lack high-resolution information about hexamers and pentamers in a global, lattice-wide context. Recent advances in cryo-electron microscopy (cryo-EM) and cryo-electron tomography (cryo-ET) have yielded additional important insights into capsid structure (9, 4, 5, 22, 23), but resolution has generally remained limited, particularly for the pentamer.

Consequently, important questions about capsid structure and assembly remain unanswered. For instance, the polyanionic HIV-1 assembly cofactor inositol hexakisphosphate (IP6) is known to coordinate a ring of electropositive charges in the central pores of Gag and CA hexamers (24–29), but the role it plays in CA pentamer structure, if any, remains uncertain. Polyanionic deoxyribonucleotide triphosphates (dNTPs) also interact with and translocate through hexamer central pores (30–32)—an observation with important implications for capsid nucleotide import and reverse transcription—but whether dNTPs, like IP6, also have a role in stabilizing capsid structure is unknown.

In this study, we developed an *in vitro* lattice ‘templating’ assembly technique that enabled us to enrich for CA pentamer formation and to use SPA to examine the lattice under a range of assembly conditions. Using this approach, we demonstrated that pentamer formation is strictly dependent upon polyanion coordination, providing a key advance in our understanding of capsid assembly and, more broadly, HIV-1 maturation. We also used lattice templating to determine structures of the lattice bound to the first-in-class capsid-targeting antiretroviral compound GS-6207 (Lenacapvir). We found that GS-6207 binds exclusively to hexamers, a behavior that arises from key structural differences between the two oligomers. Altogether, this work provides new insights into the molecular mechanisms of HIV-1 capsid assembly, structure, and function.

## RESULTS

### Structure of HIV-1 CLPs determined via cryo-ET

Although a good general understanding of HIV-1 capsid structure has been available for some years, high-resolution structures of HIV-1 hexamers and pentamers in the context of the mature lattice have not been determined, which limits the understanding of the principles of capsid structure and assembly. We therefore sought to determine high-resolution structures of these complexes using cryo-electron tomography (cryo-ET) with subtomogram averaging and single particle analysis (SPA) cryo-electron microscopy (cryo-EM). As reported previously (25), purified wild-type HIV-1 CA assembles efficiently into capsid-like particles (CLPs) at pH 6.2 in presence of the essential assembly cofactor inositol hexakisphosphate (IP6) (Figure 1A, Supplemental Figure 1B), which is coordinated within the CA hexamer pore by residues R18 and K25 in α-helix 1 (Supplemental Figure 1A). These CLPs have a predominantly conical morphology resembling that of authentic HIV-1 capsids (33, 3, 34, 5). The hexamer pore has also been proposed to function as a deoxyribonucleotide triphosphate (dNTP) import channel (30, 32). We therefore also assessed whether dNTPs stimulate CLP assembly but found that they do not.

We imaged IP6-assembled CLPs by cryo-ET (Supplemental Table 1, Figure 1B-D) and determined the structures of their constituent hexamers by subtomogram averaging to an estimated resolution of 3.9 Å. Using the aligned hexamer subtomogram positions, we constructed lattice maps that mark the positions of hexamers and pentamers within the CLPs (Figure 1B). As in authentic HIV-1 capsids (3–5), the conical CLPs consist of a lattice of CA hexamers closed by 12 pentamers. Finally, we applied subtomogram averaging to pentamer positions identified in the lattice maps, resulting in a pentamer structure with an estimated resolution of 6.2 Å (Figure 1D).

The structures of both hexamers and pentamers in the CLPs are the same as those previously determined at lower resolution from capsids in authentic HIV-1 viral particles (5). We observed strong densities corresponding to established and hypothesized IP6 binding sites in the central pores of both hexamers and pentamers (Figure 1C,D) (25, 26, 29), in agreement with a recent low-resolution cryo-ET study of authentic virions (23). While the hexamer structure in published crystal structures is equivalent to that of hexamers within authentic capsids (Figure 1C and (5, 18)), the structure of the pentamer within authentic capsids differs from crystal structures of crosslinked pentamers (Supplemental Figure 1C and (17), raising the question of whether the internal environment of the virion or other viral components are required to stabilize the ‘viral’ pentamer. Our data demonstrate that a minimal system consisting of only CA and IP6 is sufficient to reconstitute the authentic in-virus structures of both the hexamer and the pentamer, as well as the conical fullerene architecture of the capsid.

### Structure of the HIV-1 CLP lattice determined via SPA cryo-EM

We next sought to generate higher resolution structures of the CLP pentamer using single particle analysis (SPA) cryo-EM (Supplemental Table 1). Lattice details in flat (hexamer-rich), curved (pentamer-containing), and edge regions were clearly observable in micrographs and in 2D class averages (Figure 1E). We focused our analysis on regions of the lattice containing pentamers and determined the structure of the pentamer surrounded by five hexamers (the ‘global’ pentamer structure) to an estimated resolution of 3.6 Å (Supplemental Figure 2; Figure 1F-H). In accordance with our cryo-ET data (Figure 1C,D), strong densities were present at proposed binding sites in the pentamer and at known IP6 binding sites in the adjacent hexamer (25, 26, 29). Importantly, these SPA hexamer and pentamer structures are indistinguishable from those determined via cryo-ET (Supplemental Figure 1D-E) and via an independent SPA study described in the accompanying manuscript (35), demonstrating that SPA is suitable for structural studies of complex CA lattices containing both hexamers and pentamers.

**FIGURE 2.**
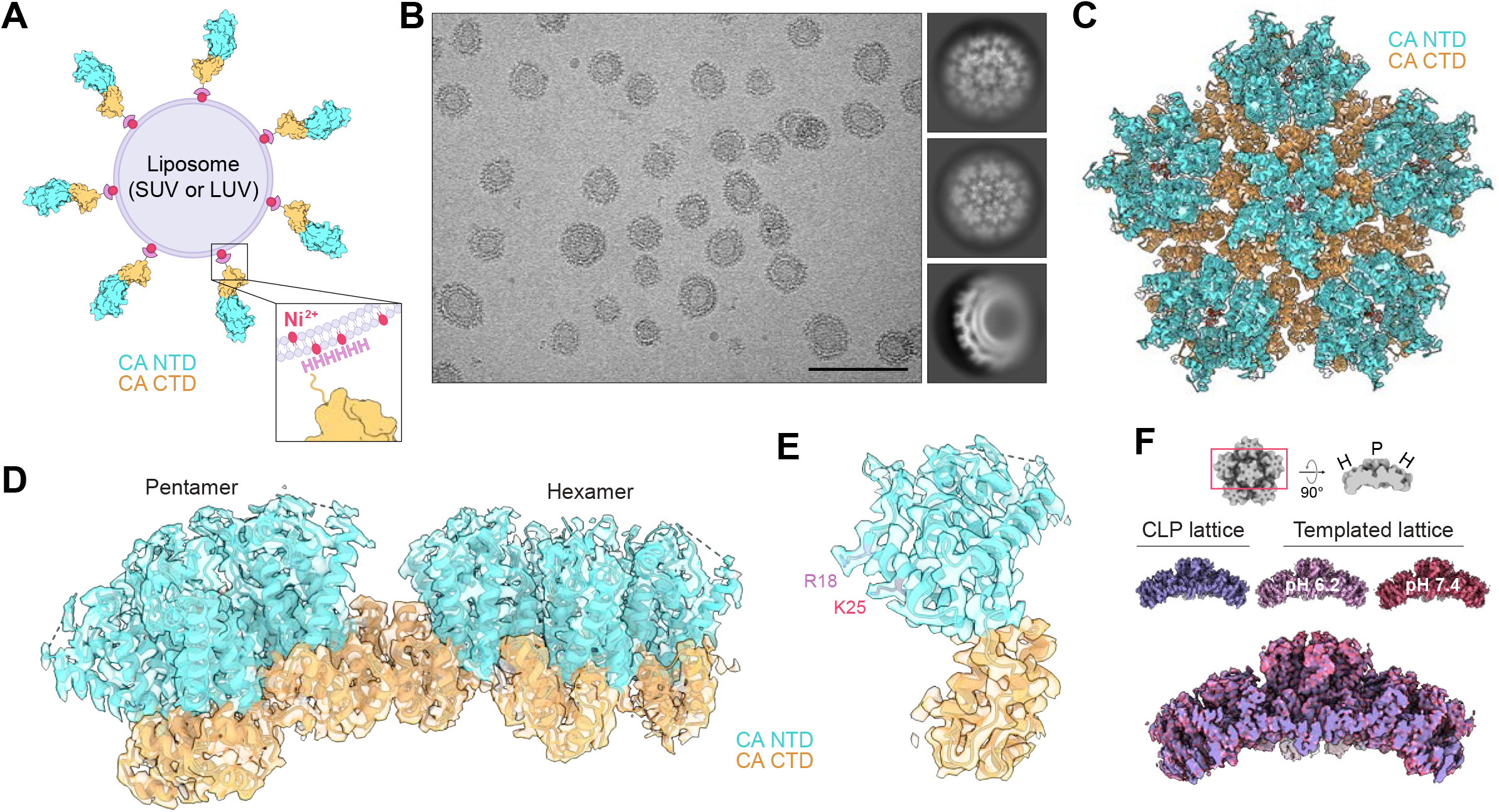
Single particle analysis of liposome-templated HIV-1 CA lattice. **A**. Diagram of CA lattice templating scheme. Purified CA-6xHis is anchored to liposomes containing nickel-chelated lipids. If applicable, polyanions are then added. SUV, small unilamellar vesicle; LUV, large unilamellar vesicle. **B**. Cryo-EM micrograph of SUV-templated CA with IP6 (pH 7.4) (left) with select 2D class averages (right). Scale bar, 100 nm. **C-E**. Atomic model for liposome-templated CA lattice (prepared at pH 7.4 in the presence of IP6) with corresponding cryo-EM map. *myo*-IP6 molecules are docked according to best cross-correlation with the cryo-EM density. The CA NTD is shown in cyan and the CA CTD is shown in orange. C, ‘global’ pentamer structure (pentamer surrounded by its five nearest hexamer neighbors). D, focused view of adjacent pentamer and hexamer. E, focused view of a single CA pentamer monomer with IP6-coordinating R18 and K25 sidechains highlighted in lavender and magenta. **F**. Top: diagram of map side views shown beneath. ‘H,’ hexamer; ‘P,’ pentamer. Middle: side views of ‘global’ pentamer cryo-EM maps for capsid-like particle (CLP) lattice (purple) and SUV-templated lattice prepared at pH 6.2 or pH 7.4. All were assembled with IP6. Bottom: superimposed maps. Corresponding atomic models are shown in Supplemental Figure 3E.

### HIV-1 CA lattice formation via protein templating on lipid vesicles

As in isolated authentic capsids (3–5), pentamers in our CLPs were quite scarce, present at a pentamer to hexamer ratio of ∼1:20. We therefore wondered whether the resolution of the pentamer map could be further improved by increasing this ratio. Given the relationship between pentamers and lattice curvature, we speculated that pentamers could be enriched by forcing CA to form a high-curvature lattice similar to that permitted by pentamers at the ends of authentic capsid cores (3–5). Indeed, we and others have shown that CA from other retroviruses can be induced to assemble into highly-curved icosahedral particles enriched in pentamers under specific buffer and assembly conditions (36–38). For HIV-1, however, this is possible only with the introduction of mutations (39, 40).

We therefore devised a lattice assembly scheme in which purified wild-type HIV-1 CA-6xHis is anchored to highly curved ∼30 nm small unilamellar vesicle (SUV) scaffolds doped with nickel-chelated lipids (Figure 2A). Importantly, the extreme C terminus of the CA CTD (G222-L231) is not known to make critical contacts for mature lattice formation, as this region is disordered past V221 in cryo-EM and crystal structures (5, 18). Curiously, we found that this ‘templated’ CA forms a lattice that partially covers the SUVs independently of IP6 at both pH 6.2 and pH 7.4, while addition of either IP6 or dNTPs drives full assembly of the lattice around the SUVs (Supplemental Figure 3A). By visual inspection, the assembled lattice does not differ between the two pH conditions or in the presence or absence of either IP6 or dNTPs, and the lattice generally follows the curvature of the SUVs. We also tested whether a lattice could form on larger ∼100-nm large unilamellar vesicles (LUVs) in the presence of IP6 and found that it indeed does, but it also appears to deform the liposome membrane at sites of pentamer incorporation (Supplemental Figure 3B,C). These results indicate that CA can be induced to form a highly curved lattice, and that unlike CLP assembly, formation of the templated lattice is much less dependent on pH or the presence of IP6.

**FIGURE 3.**
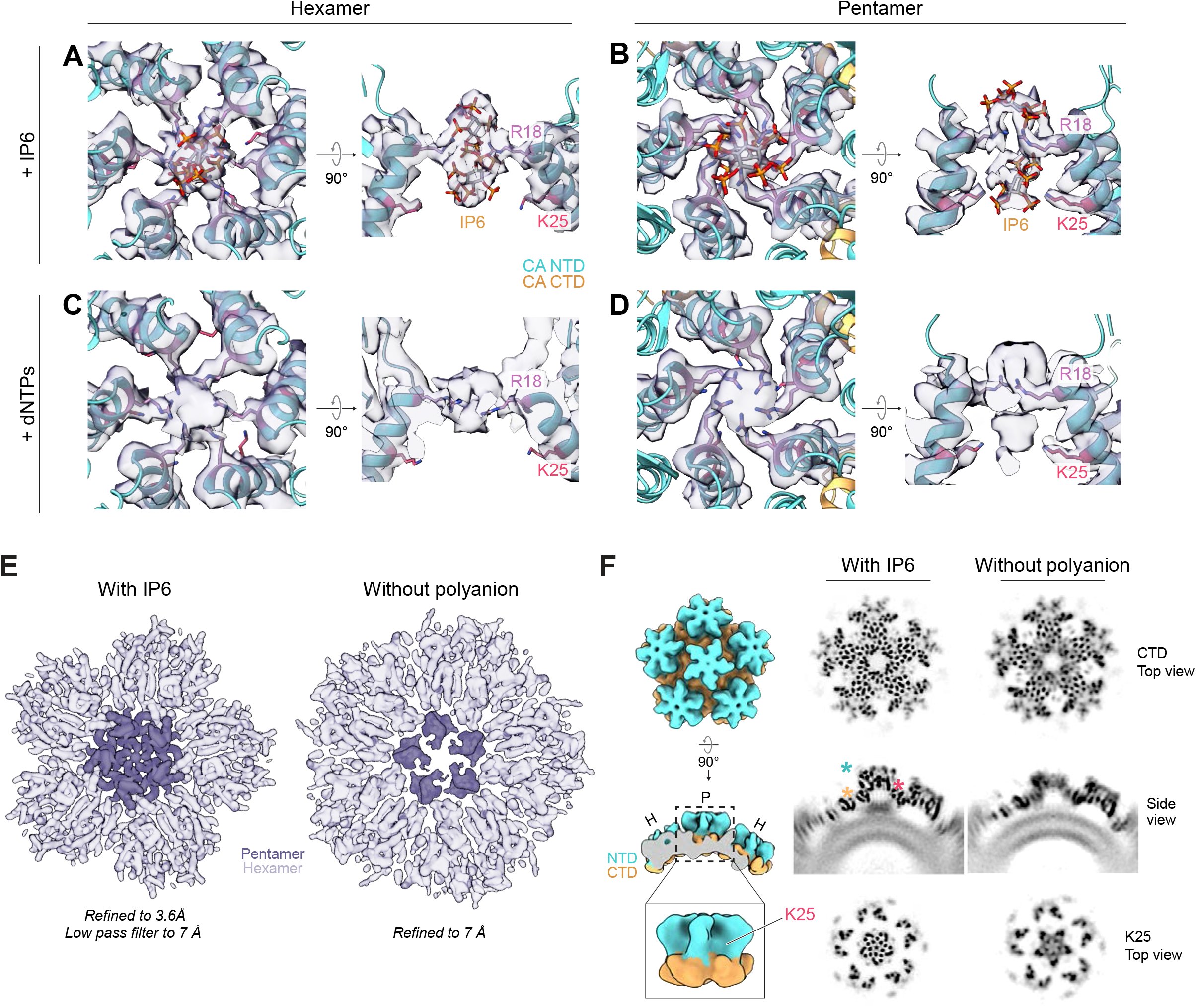
Polyanions are required for HIV-1 CA pentamer formation. **A-D**. Atomic models and corresponding cryo-EM densities of polyanions coordinated in CA hexamer (A,C) and pentamer (B,D) central pores. Lattice was prepared via liposome templating of CA-6xHis at pH 7.4 in the presence of IP6 (A-B) or dNTPs (C-D). In A-B, *myo*-IP6 molecules are docked according to best cross-correlation with the cryo-EM density. For C-D, hypothesized dNTP orientations are shown in Supplemental Figure 4A. In all A-D side views, helices from only two CA monomers are shown for clarity. **E**. Left, ‘global’ cryo-EM maps of templated HIV-1 CA-6xHis lattice prepared at pH 7.4 in the presence or absence of IP6. Pentamer (left) and aberrant pentamer (right) regions are highlighted with a darker color. To ensure a fair comparison with the “Assembled without polyanion” map (right), the “Assembled with IP6” map (left) was prepared from a subset of ∼11,000 particles randomly selected from the dataset used to determine the structure presented in Figure 2 and was low pass filtered to 7 Å. **F**. Left: diagram of regions of interest shown in orthoslice views at right. P, pentamer; H, hexamer. Right: orthoslice side and top views of the ‘global’ cryo-EM maps shown in E. Top views show the region containing pentamer K25 or pentamer CTDs. Asterisks mark positions of CA residue K25 (magenta), the CA NTD (cyan), and the CA CTD (orange). For the models in A-D and diagram in F, the CA NTD is shown in cyan and the CA CTD is shown in orange.

### Structures of templated CA lattice at pH 6.2 and at pH 7.4 determined via SPA cryo-EM

To determine whether templated CA lattice is comparable to that of CLPs and authentic capsids, we determined the structures of SUV-templated CA lattice in the presence of IP6 at pH 6.2 (Supplemental Figure 4A) and at pH 7.4 (Figure 2B-E and Supplemental Figure 4A) by SPA (Supplemental Figure 2). The estimated resolutions for the global pentamer structures were 3.3 Å and 3.1 Å, respectively—both significantly higher than that of the CLP global pentamer (Figure 1H). We noted a previously-reported (30) pH-dependent shift in the position of the hexamer CA NTD β hairpin situated above the central pore (Supplemental Figure 4B). As predicted, the increased lattice curvature results in a significant increase in the pentamer to hexamer ratio, from ∼1:20 to ∼1:5 (Supplemental Figure 3D), suggesting that fewer hexamers are needed to form a complete lattice around the highly-curved SUVs. This is similar to highly-curved regions of authentic capsids, which contain fewer hexamers per pentamer than do flatter regions of the lattice (5). By comparison, the pentamer to hexamer ratio in LUV-templated lattice is ∼1:20, reminiscent of the ratio found in our CLPs and in authentic virions (3–5).

**FIGURE 4.**
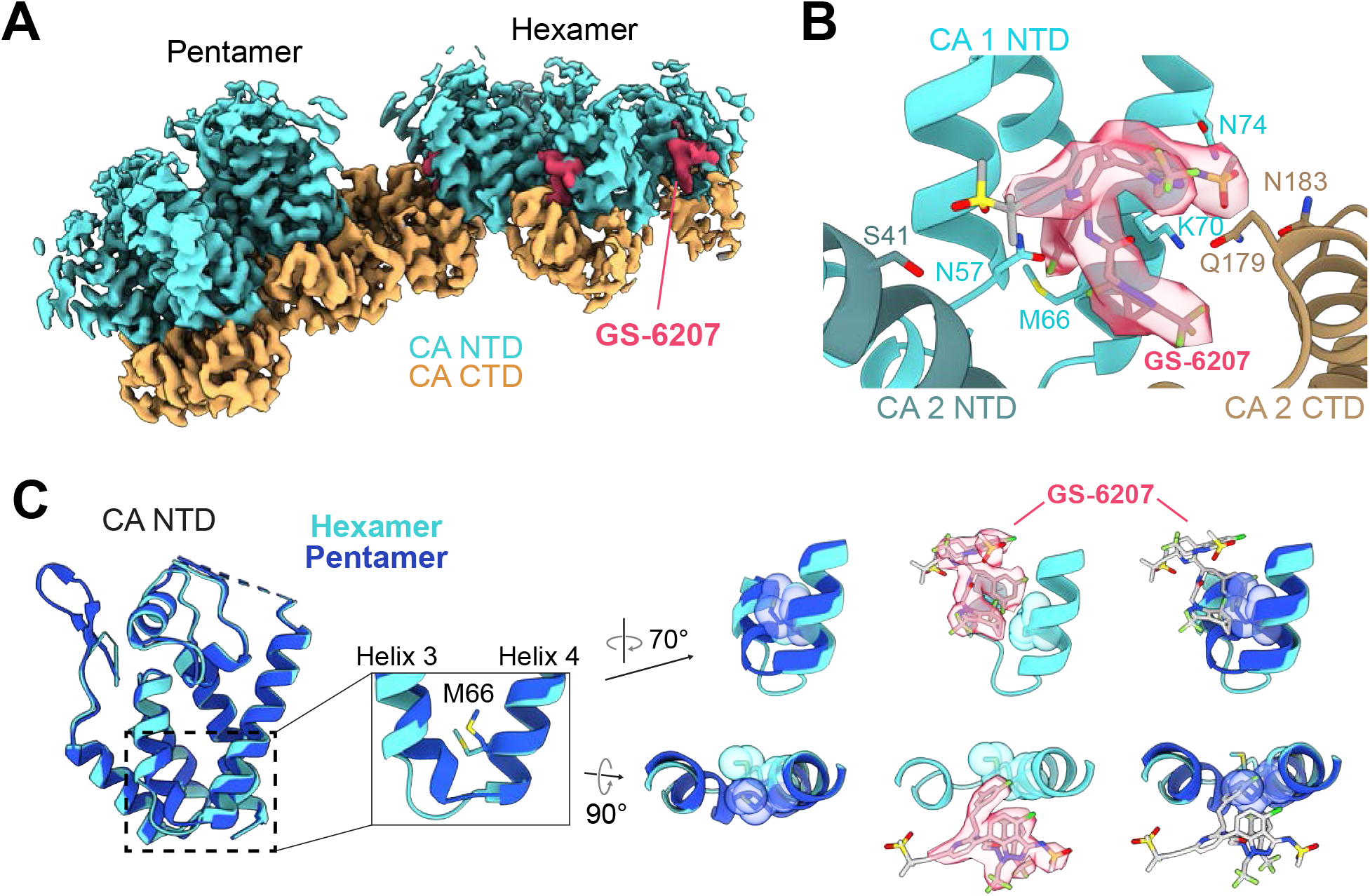
Lattice templating reveals new insights into GS-6207 binding. **A**. Cryo-EM map of GS-6207 bound to the CA lattice. Lattice was prepared via SUV templating of CA-6xHis at pH 7.4 in the presence of IP6. GS-6207 density is shown in magenta. **B**. Atomic model of the GS-6207 binding pocket between adjacent hexamer CA monomers for the structure shown in A. GS-6207 is docked according to best cross-correlation with the cryo-EM density. CA residues that have been reported to interact with GS-6207 are labeled. **C**. Comparison of hexamer (cyan) and pentamer (blue) CA NTDs from the structure shown in Left: overlaid hexamer and pentamer NTDs. Middle: M66 ‘gating’ residue in the hexamer-pentamer ‘switch’ region between ɑ-helices 3 and 4. Right: View of the hexamer CA NTD GS-6207 binding site compared with the same region for the pentamer. GS-6207 is docked according to best cross-correlation with the cryo-EM density. For the map and model shown in A-B, the CA NTD is shown in cyan and the CA CTD is shown in orange.

The global pentamer structures are virtually indistinguishable between CLP and templated lattice preparations (Supplemental Figure 4A). This observation, along with the ability of templated CA to accommodate a broader range of assembly conditions, indicates that liposome templating is a useful tool for structural studies of the lattice under more physiological buffer conditions than have previously been feasible for HIV-1 CA. In addition, the curvature imposed on the CA lattice by templating on SUVs or LUVs does not alter the native geometry of the CA pentamer or its nearest hexamer neighbors, highlighting the strict local lattice curvature imposed by the pentamer (Figure 2F, Supplemental Figure 3C; see also accompanying manuscript (35).

### Pentamer formation is dependent upon polyanion coordination

We next sought to understand the role of IP6 in CA lattice formation. In agreement with the CLP data (Figure 1C,D,G), we noted strong densities corresponding to two established IP6 binding sites in hexamer central pores from the SUV-templated CA lattice prepared at pH 7.4 (25, 26): a primary binding site above the R18 ring, and a secondary site near K25 below the ring (Figure 3A). We also observed two densities at similar positions within the central pore of the pentamer, although the contribution of K25 to IP6 coordination appeared stronger in the pentamer than in the hexamer (Figure 2E, Figure 3B). We noted the same features in lattice assembled in the presence of dNTPs instead of IP6 (Figure 3C,D) (global pentamer map estimated resolution of 3.5 Å; Supplemental Figure 2), indicating that both polyanions can drive formation of hexamers and pentamers under these conditions. We were unable to determine dNTP orientation or number within either oligomer, but we suggest some possibilities based on published molecular dynamics simulations (32) in Supplemental Figure 5A.

Next, we compared the cryo-EM dataset used to generate the IP6-containing CA lattice structure presented in Figure 2 and in Figure 3A-B to a dataset containing SUV-templated CA in the absence of any polyanion (Figure 3E,F; Supplemental Figure 5B). Intriguingly, while hexamers assembled normally in the absence of polyanion, pentamers were severely defective. They were also rarer than in lattice assembled with polyanions: the ratio of these aberrant ‘pseudo-pentamers’ to hexamers was ∼1:10, in contrast to the 1:5 ratio seen in templated lattice assembled with polyanion. We generated a pseudo-pentamer global structure refined to an estimated resolution of 7.0 Å and found that the CTDs of CA monomers appear fairly normal, while the NTDs are largely unresolved (Supplemental Figure 5C). These data suggest that pentamer formation is strictly dependent on coordination of polyanions such as IP6, while hexamer formation is not under these conditions. Indeed, hexamer pores in our no-polyanion templated lattice dataset contain a density much smaller than that that of IP6 (Supplemental Figure 5B), which we postulate corresponds to a small anion that helps neutralize the R18 charge (41), allowing hexamers to form.

To assess the importance of polyanion coordination *in vivo*, we generated several R18 (Gag R150) and K25 (Gag K157) CA mutants and measured their infectivity and release in tissue culture (Supplemental Figure 5E-G). In agreement with published studies (30, 25, 42, 29), all mutants had drastically reduced infectivity, likely due to decreased virus-like particle (VLP) release from infected cells. Mutations to K25 have previously been shown to also have deleterious effects on capsid morphology and stability (28, 29), so we also examined the capacity of purified CA_K25N_ to assemble in the presence of IP6 (Supplemental Figure 5D). Intriguingly, we found that it forms mostly tubular assemblies, which are known to contain only hexamers (4, 9). This result is consistent with the tendency of purified HIV-1 CA to form tubes, but not polyhedrons, in the absence of IP6 under high-salt (1 M NaCl) conditions (10, 43, 44). We propose that a major role of IP6 in HIV-1 virion production is to drive pentamer formation and, thus, complete assembly of the capsid around the viral genome.

### Lenacapavir selectively binds to the CA hexamer

Finally, we extended our templating approach to understanding the interaction of the capsid-targeting antiretroviral compound GS-6207 (marketed by Gilead Sciences as Lenacapavir) with CA. We determined the structure of this compound bound to templated CA lattice at an estimated resolution of 3.1 Å. Consistent with crystal structures of crosslinked CA hexamers bound to GS-6207 (45, 46), we found that the compound binds to the ‘FG binding pocket’ between the NTD and CTD of adjacent hexamer monomers (Figure 4A-B, Supplemental Figure 2). Importantly, this site is also targeted by FG motif-containing host cell proteins CPSF6, Nup153, and Sec24C (19, 20, 47, 45, 46, 21, 35). GS-6207 density was completely absent from the pentamer, indicating that the compound binds exclusively to hexamers. This selectivity appears to arise from spatial constraints imposed by subtle conformational differences between pentameric and hexameric CA, as the FG binding pocket is altered in the pentamer by a structural ‘switch’ between NTD ɑ-helices 3 and 4. (For more detailed structural comparisons, see (35) and (40).) This switch alters the position of M66 in the pentamer such that it would sterically interfere with the difluorobenzyl group of GS-6207 (Figure 4C), providing an explanation for the selective binding of GS-6207 to hexamers.

## DISCUSSION

Application of cryo-electron tomography (cryo-ET) with subtomogram averaging to our *in vitro* capsid-like particle (CLP) assembly system confirmed that the distributions and structures of hexamers and pentamers in CLPs match those previously described at lower resolution for authentic HIV-1 capsids (33, 34, 5, 23). Single particle analysis (SPA) cryo-electron microscopy (cryo-EM) of these CLPs (see also accompanying manuscript (35) and preprint from (40)) enabled us to collect larger datasets than feasible with cryo-ET, resulting in higher-resolution structures. This was particularly advantageous for the rarer pentamer. The structure of the pentamer within the capsid/CLP lattice differs significantly from published crystal structures of crosslinked pentamers (17), and instead matches the structure of the pentamer within the virion. This indicates that a combination of a curved hexameric lattice framework and IP6 is sufficient to reconstitute the viral pentamer structure—no other virus-specific factor is required.

Although our CLP assembly scheme recapitulates the morphology and composition of authentic capsids, it only works at low-pH (pH 6.0-6.2) conditions. This is not ideal for studies of capsid interactions with host cell factors and potential capsid-targeting antiretroviral compounds, which, in nature, would interact with the capsid at a physiological pH. The liposome scaffold-based lattice templating technique we introduce here enabled us to overcome this limitation and to study lattice assembled under more physiological buffer conditions. A second advantage of the lattice templating system is that it itself promotes lattice formation, likely by simply increasing proximity of capsid protein (CA) monomers: we found that templated CA readily forms some degree of lattice in the absence of typical drivers of *in vitro* CA lattice assembly such as the essential cofactor inositol hexakisphosphate (IP6), which promotes conical CLP and capsid formation (25, 29), or high concentrations of salt (1 M NaCl), which promotes formation of CA tubes composed exclusively of hexamers (43, 44, 10, 9, 4).

Close inspection of such lattice by cryo-EM revealed that CA hexamers can form normally under these conditions, but pentamers cannot. Our structural data indicate that K25 is particularly important for IP6 coordination in the pentamer, and we found that purified mutant CA_K25N_ cannot assemble into conical CLPs, even in the presence of IP6, but instead forms hexamer-rich tubes. This suggests that polyanions—specifically IP6, given its established importance for virus production and capsid stabilization—are indispensable for pentamer formation. In recent years, IP6 has been shown to be important also for lentiviruses EIAV, SIV, FIV, and BIV (42, 48, 27) and the alpharetrovirus RSV (38). Given this, and the conservation of regularly spaced Arg or Lys residues in ɑ-helix 1 of CA (49), it we hypothesize that the reported polyanion requirement for pentamer formation for HIV-1 is conserved for other retroviruses as well.

A recent molecular dynamics simulation analysis of capsid assembly (50) lends further support to this model, suggesting that IP6 has a higher affinity for pentameric CA than for hexameric CA within capsids and that IP6 is critical for assembly of a high-curvature lattice. Moreover, a cryo-ET analysis (23) of capsid cores from native virions reported that the central pores of viral pentamers contained two strong densities that were predicted to be IP6. Hexamer pores contained much weaker density, although exposure of the capsid cores to excess IP6 yielded hexamer structures with clearer densities. The number and mobility of IP6 molecules coordinated by capsid CA in nature remains an area of active research (32, 51, 23, 50), but our work suggests that IP6 coordination within the pentamer is likely quite stable. We also found that dNTPs can substitute for IP6 to drive full assembly of SUV-templated CA lattice. Capsid hexamer pores are thought to facilitate nucleotide import for reverse transcription (30, 32), but whether pentamer pores can also import nucleotides is unknown. Furthermore, whether nucleotides, like IP6, have a role in physiological capsid stabilization remains to be determined.

Lastly, we determined the structure of templated lattice bound to the potent antiretroviral compound GS-6207 and found that the compound binds exclusively to hexamers at the ‘FG’ binding site between neighboring CA monomers. We attribute this selective binding to a structural ‘switch’ in the CA NTD that promotes a conformational change in CA monomers and drives oligomerization to either the pentameric or the hexameric state ((40) and (35)). This conformational change reshapes the FG binding pocket in pentamers, resulting in a steric block of GS-6207 access to the site. Work described in the accompanying manuscript demonstrates how this remodeling also prevents binding of FG motif-containing host cell proteins CPSF6, Nup153, and Sec24C to pentamers (35). These findings have important implications for understanding how the HIV-1 capsid interacts with host cell proteins and for ongoing efforts to pharmacologically target the capsid.

Altogether, our work provides key new insights into HIV-1 capsid structure and assembly and demonstrates that SPA cryo-EM is a valuable tool for high-resolution structural virology. In addition, the lattice templating method we introduce here has proven to be a valuable platform for interrogating the molecular mechanisms underlying lattice assembly and lattice interactions with small molecules and proteins of interest. We expect that this technique can be extended to studies of other viruses as well.

## MATERIALS AND METHODS

### Protein expression and purification

Both untagged HIV-1 CA and HIV-1 CA-_6x_His were produced in BL21 *Escherichia coli* cells. Cells were grown to OD_600_ ∼0.8 before induction with 400 uM Isopropyl β-d-1-thiogalactopyranoside (IPTG) at 30°C for 6 hr, then collected by centrifugation. Cells were resuspended in Lysis Buffer (for untagged CA, 25 mM Tris pH 8.0, 2 mM phenylmethylsulfonyl fluoride (PMSF), 4 mM tris(2-carboxyethyl)phosphine (TCEP); for CA-_6x_His, 50 mM Tris pH 8.0, 300 mM NaCl, 10 mM imidazole, 2 mM PMSF, 4 mM TCEP) and lysed by sonication. Insoluble material was collected by ultracentrifugation and discarded. Polyethylenimine (PEI) was added to the cleared lysate to 0.3% and mixed at 4°C for 10 min. Bulk protein was then precipitated from the lysate with 20% ammonium sulfate (incubated, stirring, at 4°C for 40 min), and precipitated protein was collected by centrifugation. The protein pellet was resuspended in CA Lysis Buffer, then run though a HiPrep Desalting column (GE Healthcare Life Sciences) with 50 mM Tris pH 8.0, 2 mM TCEP. Protein peak fractions were collected and combined.

For purification of untagged CA and CA_K25N_, CA was isolated from the desalted crude protein via a tandem HiTrap Q HP-HiTrap SP HP anion-cation exchange column setup: crude protein was applied to the columns (Q first, followed immediately by SP), and flow-through containing CA was collected. Residual protein was collected by washing the column once with 25 mM Tris pH 8, 2 mM TCEP, then twice with 25 mM Tris pH 8, 25 mM NaCl, 2 mM TCEP. Eluate and washes were combined and buffer exchanged into 25 mM Tris pH 8, 2 mM TCEP via gel filtration (Superdex 75 Increase 10/300; GE Healthcare Life Sciences). Protein peak fractions were combined and concentrated to 1 mM, then flash-frozen in liquid nitrogen and stored at -80°C.

For purification of CA-_6x_His, NaCl and imidazole were added to the desalted crude protein to final concentrations of 300 mM and 10 mM, respectively. The solution was then incubated with NiNTA agarose resin (Qiagen) for 1 hr at 4°C, rotating. The resin was washed 1x in batch with 20 mL CA-His Lysis Buffer, then transferred to a gravity column and washed 10x with 1 mL CA-His Wash Buffer (50 mM Tris pH 8.0, 300 mM NaCl, 30 mM imidazole, 2 mM TCEP). CA-_6x_His was then eluted in 1 mL fractions with CA-His Elution Buffer (50 mM Tris pH 8.0, 300 mM NaCl, 300 mM imidazole, 2 mM TCEP). CA-_6x_His was further purified by gel filtration (Superdex 75 Increase 10/300; GE Healthcare Life Sciences) in CA-His Gel Filtration Buffer (25 mM Tris pH 8.0, 2 mM TCEP). Protein peak fractions were combined and concentrated to 500 uM, then flash-frozen in liquid nitrogen and stored at -80°C.

### *In vitro* CA capsid-like particle (CLP) assembly

CLP assembly reactions were buffered with 50 mM MES pH 6.2 and contained 500 uM purified untagged HIV-1 CA, 2.5 mM inositol hexakisphosphate (IP6) (Tokyo Chemical Company), and 1 mM TCEP. CA was first warmed at 37°C for 5 min, then combined with IP6 and incubated at 37°C for 15 min. Assemblies were stored on ice or at 4°C until grid preparation. Templated assemblies were screened by negative stain transmission electron microscopy (TEM) (FEI Tecnai 12 BioTwin TEM or FEI Morgagni TEM; see Negative stain EM methods section) before cryo grid preparation.

### Liposome preparation

Chloroform stock solutions of 1,2-dioleoyl-sn-glycero-3-phosphocholine (DOPC) and 1,2-dioleoyl-sn-glycero-3-[(N-(5-amino-1-carboxypentyl)iminodiacetic acid)succinyl], nickel salt (DGS-NiNTA) were purchased from Avanti Polar Lipids. Cholesterol was purchased from Nu-Chek Prep (Elysian, MN), and cholesterol stock solutions were prepared by standard gravimetric procedures to within 0.2% error. Stock solutions were mixed at an 85:10:5 DOPC:DGS-NiNTA:cholesterol ratio, then exchanged into 50 mM MES with 2 mM TCEP (pH 6.2 or 7.4) via rapid solvent exchange to a final total lipid concentration of 13 mM. SUVs were prepared by sonication in a room temperature water bath for 30 min (Sonblaster Series 200, Narda Ultrasonics Corporation), then filtered through a desalting spin column (Thermo Fisher Scientific). LUVs were prepared by extrusion through a 100-nm polycarbonate filter using a mini-extruder (Avanti Polar Lipids) 25 times at room temperature.

### CA templating on liposomes

Purified HIV-1 CA-_6x_His was combined with liposomes and incubated at 37°C for 5 min. 10 mM IP6 (Tokyo Chemical Company) in 50 mM MES (pH 6.2 or 7.4) was added to a final concentration of 1 mM, and the assembly reaction was incubated at 37°C for 15 min. SUV assembly reactions (pH 6.2 or 7.4) contained 330 uM CA-_6x_His and 5.9 uM SUVs; LUV assembly reactions (pH 7.4) contained 190 mM CA-_6x_His and 3.7 mM LUVs. If applicable, dNTPs (equimolar mixture of dATP, dCTP, dGTP, dTTP; Cytiva) were added to a final concentration of 2 mM in place of IP6. For preparation of GS-6207-bound lattice, GS-6207 (Gilead Sciences) was added to a final concentration of 330 uM 5 min after IP6 addition, and the reaction contained 125 mM NaCl and was buffered with 50 mM HEPES (pH 7.4) rather than MES. Templated assemblies were screened by negative stain TEM (FEI Tecnai 12 BioTwin TEM; see Negative stain EM methods section) before cryo grid preparation.

### Virus production and infectivity assay

HIV-1_ΔEnv_ consisted of NL4-3-derived proviral vector with a 5’ cytomegalovirus (CMV) driven green fluorescent protein (GFP) and defective for Vif, Vpr, Nef, & Env (kindly provided by Vineet Kewal-Ramani, National Cancer Institute-Frederick). CA mutations were made using the In-Fusion cloning system (Takara Bio) using either custom Gene Blocks (Integrated DNA Technologies) (R18A, R18L, R18K, K25A, K25E, K25R, K25N, K25N-N21K) or PCR site-directed mutagenesis (PR-D25A). All plasmids were verified by sequencing. The plasmid for Vesicular Stomatitis Virus glycoprotein (VSV-g, NIH AIDS Reagent Program) has been previously described (52).

HIV-1 virus-like particles (VLPs) were produced by Lipofectamine 3000 (Thermo Fisher Scientific) transfection of 293FT cells (purchased from Invitrogen; cells maintained as previously described in (27)) at ∼50-60% confluence with 900 ng of the proviral vector and 100 ng of VSV-g. Media containing VLPs (‘viral media’) was collected two days post transfection. Viral media was then frozen at -80°C for a minimum of 1 hr to lyse cells, briefly thawed in a 37°C water bath, then precleared by centrifugation at 3000 x g for 5 min. The supernatant was collected and added to fresh HEK293FT cells (6-well format) at low multiplicity of infection to prevent infection saturation. Infected cells were collected, washed with phosphate-buffered saline (PBS; 137 mM NaCl, 2.7 mM KCl, 10 mM Na_2_HPO_4_, 1.8 mM KH_2_PO_4_), and treated with 10 mM TrypLE™ Express Enzyme (Gibco/Thermo Fisher Scientific). Cells were then resuspended in PBS, and 10% PFA was added to a final concentration of 5%. After 10-20 min incubation at room temperature, the cells were collected by centrifugation at 500 x g for 5 min and resuspended in 500 μL of PBS. Cells were assayed for fluorescence using an Attune Flow Cytometer with proprietary Attune collection and analysis software (Thermo Fisher Scientific).

Cell lysis and preparation of released VLPs was carried out as previously described (27). VLPs were concentrated over 500 μL of 20% sucrose in PBS at 50,000 x g for 45 min (Beckman Coulter Optima TLX). Infections were assessed by western blot using the following antibodies: RbαHIV-p24 (provided by the HIV Reagent Program), AlexaFluor680-conjugated GtαRb-IgG (Life Technologies, A21076), and AlexaFluor790-conjugated MsαGAPDH (Santa Cruz Biotech, sc-365062). Blots were imaged using an Odyssey Imaging System (Li-COR), and band densities were measured using Image Studio Lite software (Li-COR). Raw band densities were imported into Microsoft Excel, and infections were normalized to percent of infections for wild-type virus, followed by normalization to a GAPDH loading control. Normalized data were then exported to CSV format for ANOVA and graph generation in Rstudio (53, 54).

### Negative stain EM of CLPs

4 µL sample was applied to a glow-discharged grid (Electron Microscopy Sciences Formvar/Carbon 200 Mesh) for 30 sec. The grid was washed with 100 uM IP6 in 50 mM MES (pH 6.2 or 7.4) for ∼5 sec, then stained with 2% uranyl acetate for 2 min and blotted dry. Samples were imaged at 120 kV (FEI Tecnai 12 BioTwin TEM) or 100 kV (FEI Morgagni TEM).

### Cryo-ET and SPA cryo-EM sample preparation

CA CLPs (3-4 uL sample) were applied to a glow-discharged grid (Protochips 2/2-3Cu C-Flat grids; PELCO easiGlow glow discharger), then blotted for 2-3 sec and plunge-frozen in liquid ethane. Plunging was performed at 90-100% humidity via automatic backside blotting (via Leica Microsystems Cryo GP 2 plunge freezer) or dual-side blotting (Mark IV FEI/Thermo Fisher Scientific Vitrobot). CLP samples for cryo-ET contained BSA-coated 10 nm colloidal gold at a CLP:gold bead ratio of ∼8:1.

Templated CA (3.5 uL sample) was applied to a glow-discharged grid (QUANTIFOIL 300 mesh Au 1.2/1.3), then manually wicked away from the edge of the grid with a Kimwipe (Kimberly-Clark). An additional 3.5 uL sample was applied, then automatically dual-side blotted for 3 sec and plunge-frozen in liquid ethane at 95% humidity/4-6°C on a Mark IV FEI/Thermo Fisher Scientific Vitrobot.

### Cryo-ET and SPA cryo-EM data collection

Cryo-ET samples were imaged at 300 kV on a Titan Krios TEM (Thermo Fisher Scientific) equipped with a K2-XP direct detector (Gatan, Inc.) and a BioQuantum post-column energy filter (Gatan Inc.) with a slit width of 20 eV. Imaging was done at a nominal magnification of 105,000x with a physical pixel size of 1.379 Å/pixel. Using SerialEM (55), dose-symmetric tilt series (56) were collected with a tilt angle range between −60° and 60°, 3° increment, and dose fractionation of each tilt image into 10 frames.

SPA samples were imaged at 200 kV on a Talos Arctica TEM (Thermo Fisher Scientific) equipped with a K3 direct detector (Gatan, Inc.) and a BioQuantum energy filter (Gatan Inc.) with a slit width of 20 eV. Imaging was done at a nominal magnification of 63,000x in super-resolution mode with a pixel size of 1.31 Å/pixel. Using SerialEM (55), a total of 50 frames were captured as movies. See Supplemental Table 1.

### Cryo-ET data processing

See Supplemental Table 1.

#### Pre-processing and tomogram reconstruction

Gain correction, motion correction and dose-reweighting were performed on the raw tilt series movies using alignframes from the IMOD package (57) The defocus and angle of astigmatism of each tilt image was estimated using ctfplotter from IMOD. Tilt series were manually aligned using gold fiducials in IMOD, and CTF-multiplied tomograms were reconstructed using novaCTF (58). The reconstructed tomograms were then serially binned by factors of 2, 4 and 8 in IMOD to produce binned tomograms for use in the initial stages of subtomogram alignment.

#### Segmentation of HIV-1 CA CLPs from tomograms

8× binned, non-CTF corrected tomograms were imported into the Ilastik software package (version 1.3.2rc1) (59) to segment the CLP cores. The vertex coordinates of the segmented volumes were used to define an isosurface for each CLP in MATLAB and an oversampled grid of points with 3.3 nm spacing was defined for subtomogram extraction along this surface using the TOM toolbox (60) with an initial estimate of the appropriate Euler angles based on the normal vector to the surface at each point.

#### Initial hexamer reference construction

An initial reference was constructed using a 4-tomogram subset of the CA CLP dataset containing 35 CLPs. This subset was chosen to evenly sample the range of defocus values across the data set. Subtomograms were extracted from 8× binned, 3D CTF corrected tomograms at the sampled positions with a box size of 706 Å^3^. The extracted subtomograms were averaged to obtain a starting reference.

#### Alignment of full hexamer data set

The initial hexamer reference constructed was used as a starting reference. Subtomograms were extracted for the entire data set and subjected to iterative alignment and averaging using subTOM (link: https://github.com/DustinMorado/subTOM) based on the TOM (60), AV3 (61) and Dynamo (62) packages. After initial alignment (5° increment, 35 Å low-pass filter, C6 symmetry imposed), duplicate and misaligned points were removed, and a further iteration of alignment was performed with a 2° increment. The aligned coordinates were scaled for re-extraction from 4× binned tomograms with box dimensions of 353 Å^3^. Two iterations of alignment were performed with a search range of 5 × 2° for all Euler angles, C6 symmetry and a 32 Å low pass filter. Points that had aligned onto the same positions during this iteration were removed. Subtomogram positions were scaled for re-extraction from 2× binned tomograms, with box dimensions of 353 Å^3^. Data sets were split into odd- and even-numbered half-sets of equal size. The averages of each half-set were generated and all further alignments were done independently on the two half-datasets. After two further alignment iterations, positions were scaled to the unbinned pixel size, subtomograms were extracted from unbinned data with box dimensions of 264.8 Å and a series of fine alignment iterations were performed with 1° angular increment and C6 symmetry. The final map resolution was measured by gold-standard Fourier shell correlation (FSC) between the two half data sets.

#### Pentamer subtomogram alignment and averaging

Pentamer subtomogram positions were obtained by identifying patterns of 5 hexamers with pairwise distance constraints that satisfy pentamer geometry (5). Pentamer subtomograms were extracted at the identified positions from 4× binned tomograms using a box size of 353 Å and averaged to generate an initial reference, which was then used as a basis for iterative alignment and averaging with a low pass filter setting of 32 Å. Misaligned pentamer subtomograms were then removed by manual inspection of the lattice maps in UCSF Chimera. Pentamer subtomogram positions were scaled for use with 2× binned data and were divided into independent odd and even half-sets. Subtomograms were iteratively aligned with C5 symmetry, re-extracted from unbinned tomograms with a box size of 264.8 Å and subjected to finer alignment with a 1° increment, C5 symmetry, and a low pass filter of 10.6 Å. The final map resolution was measured by gold-standard FSC between the two half data sets.

### Single particle cryo-EM data processing

See Supplemental Table 1.

Image processing was done in RELION 4.0 (63) maintained by SBGrid (64), and CryoSPARC (65), and motion correction and CTF estimation were done with MOTIONCOR2 (66) and GCTF (67). The initial map used for subsequent alignments was generated by aligning the CLP particles against EMDB-3465 (5) low pass filtered to 40 Å. Structures were determined using the pipeline described in Supplemental Figure 2.

### Atomic model building and refinement; data availability

For atomic modeling of the ‘global’ pentamer (central pentamer with neighboring hexamers), an initial reference model was prepared in Chimera from the crystal structure of full-length hexameric CA (PDB 4XFX). Real-space refinements of the preliminary model into the map from templated CA at pH 7.4 were carried out in Phenix, and the model was manually adjusted using Coot and the ISOLDE ChimeraX plugin (68). The resulting model was used as a preliminary model for all other modeled structures, which were also prepared by real-space refinements in Phenix and manual adjustments using Coot and ISOLDE. Model quality was evaluated using MolProbity. Cryo-EM maps will be deposited into EMDB, and atomic models will be deposited into the PDB. See Supplemental Table 1. When preparing figures containing atomic models, IP6 and GS-6207 were docked according to best cross-correlation with the cryo-EM density and are not included in the atomic models.

## ACKNOWLEDGEMENTS

This work was supported by National Institute of Allergy and Infectious Diseases under awards R01AI147890, HIVE-2 Collaborative Development Program 5U54AI150472-09, and U54 AI170855-01 to R.A.D., the UK Medical Research Council MC_UP_1201/16 and the Max Planck Society to J.A.G.B.) We thank the Cornell Institute of Biotechnology for sequencing and flow cytometry resources (RRID:SCR_021727 and RRID:SCR_021740, respectively). SPA work made use of the Cornell Center for Materials Research (CCMR) Facilities supported by the National Science Foundation under Award Number DMR-1719875. Cryo-ET data was collected with support of the EM facility at the MRC Laboratory of Molecular Biology. We thank K. Spoth and M. Silvestry Ramos (Cornell U. CCMR) for SPA data collection support, V. Vogt for constructive comments during manuscript preparation, R. Feathers and J.C. Fromme for data processing and atomic modeling advice, S. Brancato for construction of the pET3xa[HIV CA-6xHis] expression plasmid and for assistance with cryo-EM data collection for the GS-6207 dataset, and members of the R.A.D., J.A.G.B., J.C. Fromme (Cornell U.), and F. Schur (IST Austria) labs for helpful discussions.

## AUTHOR CONTRIBUTIONS

R.A.D. and J.A.G.B. designed research; C.M.H. carried out SPA cryo-EM and related sample preparation; C.M.H. and R.A.D. carried out SPA cryo-EM data processing; R.A.D. prepared samples for cryo-ET; A.T. carried out cryo-ET and related data processing with guidance from J.A.G.B.; C.L.R. performed mutational analyses; C.M.H., A.T., C.L.R., J.A.G.B., and R.A.D. analyzed and interpreted data; C.M.H. and R.A.D. prepared the figures; C.M.H., A.T., and R.A.D. wrote the manuscript with input from all authors; R.A.D. and J.A.G.B obtained funding and managed the project.

## ABBREVIATIONS USED

BIV: bovine immunodeficiency virus
BSA: bovine serum albumin
CA: capsid protein
CLP: capsid-like particle
CMV: cytomegalovirus
Cryo-EM: cryo-electron microscopy
Cryo-ET: cryo-electron tomography
CTD: C terminal domain
CTF: contrast transfer function
DGS-NiNTA: 1,2-dioleoyl-sn-glycero-3-[(N-(5-amino-1-carboxypentyl)iminodiacetic acid)succinyl], nickel salt
dNTP: deoxyribonucleotide triphosphate
DOPC: 1,2-dioleoyl-sn-glycero-3-phosphocholine
EIAV: equine infectious anemia virus
FIV: feline immunodeficiency virus
FSC: Fourier shell correlation
GFP: green fluorescent protein
GS-6207: Gilead Sciences antiretroviral compound ‘Lenacapavir’
IP6: inositol hexakisphosphate
IPTG: isopropyl β-d-1-thiogalactopyranoside
LUV: large unilamellar vesicle
NMR: nuclear magnetic resonance
NTD: N terminal domain
PBS: phosphate buffered saline
PEI: polyethylenimine
PMSF: phenylmethylsulfonyl fluoride
RSV: Rous sarcoma virus
SIV: simian immunodeficiency virus
SPA: single particle analysis
STA: subtomogram averaging
SUV: small unilamellar vesicle
TCEP: tris(2-carboxyethyl)phosphine
TEM: transmission electron microscope/microscopy
VLP: virus-like particle
VSV-g: Vesicular Stomatitis Virus glycoprotein

---

**SUPPLEMENTAL FIGURE 1.**
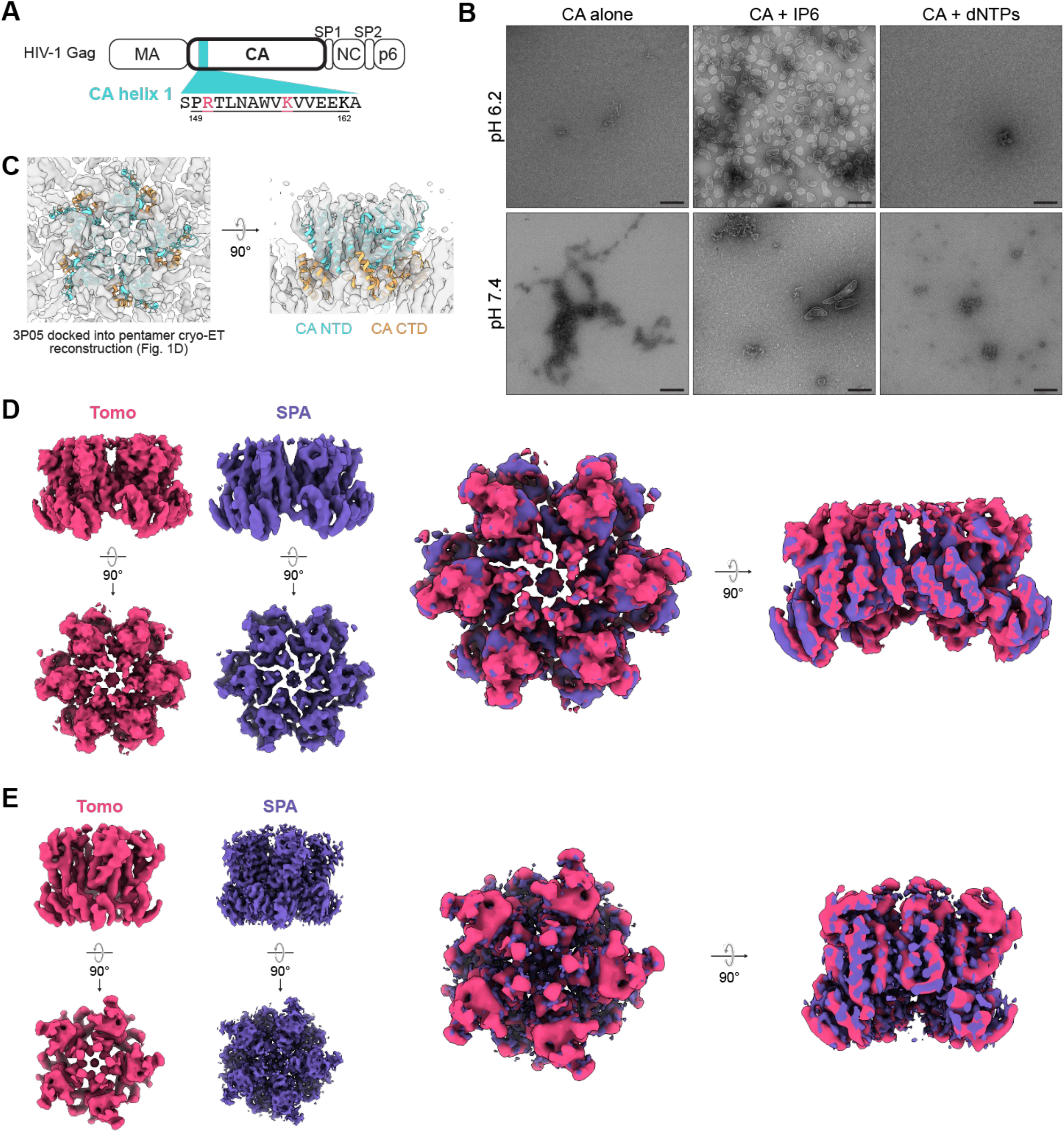
*in vitro* assembly of HIV-1 CA capsid-like particles for cryo-EM analysis. **A**. Domain diagram of full-length Gag, with CA ɑ-helix 1 highlighted in blue. Gag residues R150 and K157 (corresponding to CA residues R18 and K25, respectively) are highlighted in magenta. **B**. Negative stain TEM micrographs of *in vitro* CLP (capsid-like particle) assembly reactions containing purified CA and polyanions at pH 6.2 or 7.4. Scale bars, 100 nm. **C**. Cryo-ET reconstruction of the CLP pentamer (Figure 1D) with crystal structure 3P05 (Pornillos et al., 2011) docked according to best cross-correlation with the cryo-EM density. The CA NTD is shown in cyan and the CA CTD is shown in orange. **D-E**. Comparison of CLP hexamer (D) and pentamer (E) maps from cryo-ET (magenta) and single particle analysis cryo-EM (purple). Tomo, tomography; SPA, single particle analysis.

**SUPPLEMENTAL FIGURE 2.**
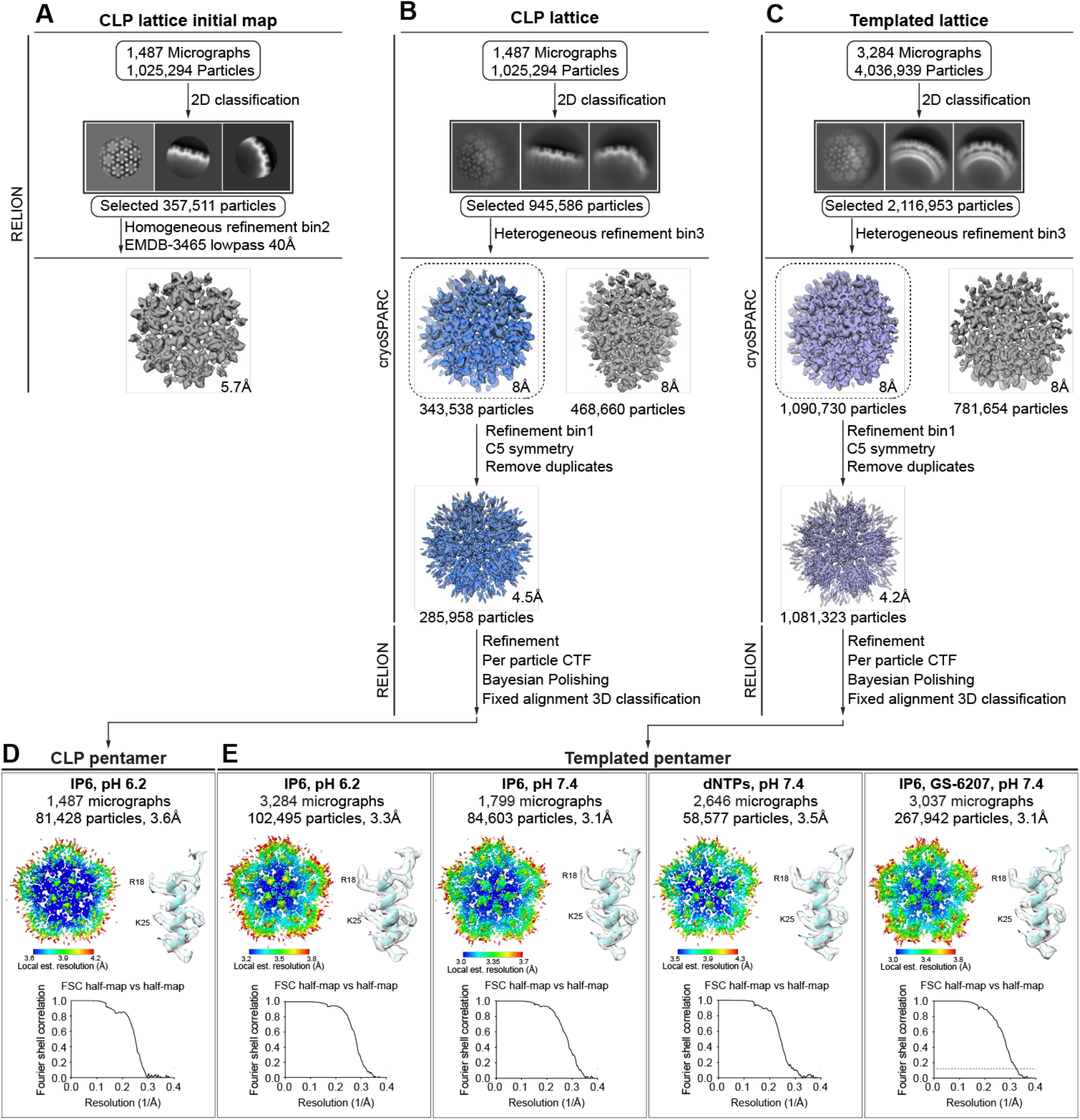
Single particle analysis data processing strategy and cryo-EM map quality. **A**. Preparation of initial reference for data processing detailed in B-C. **B-C**. Data processing strategy for ‘global’ pentamer structures (a central pentamer surrounded by its five nearest hexamer neighbors) from capsid-like particle (CLP) (B) and templated (C) CA lattice assembles. For C, the specified numbers of micrographs and initial extracted particles are for templated lattice prepared with IP6 at pH 6.2. See Supplemental Table 1 for data collection and processing details for all samples. **D-E**. Fourier shell correlation (FSC) plots and example cryo-EM densities for SPA cryo-EM maps and models from CLP (D) and liposome-templated (E) CA lattice assemblies. 0.143 FSC threshold is marked with horizontal dotted lines. Example cryo-EM densities show residues 13-31 in CA ɑ-helix 1.

**SUPPLEMENTAL FIGURE 3.**
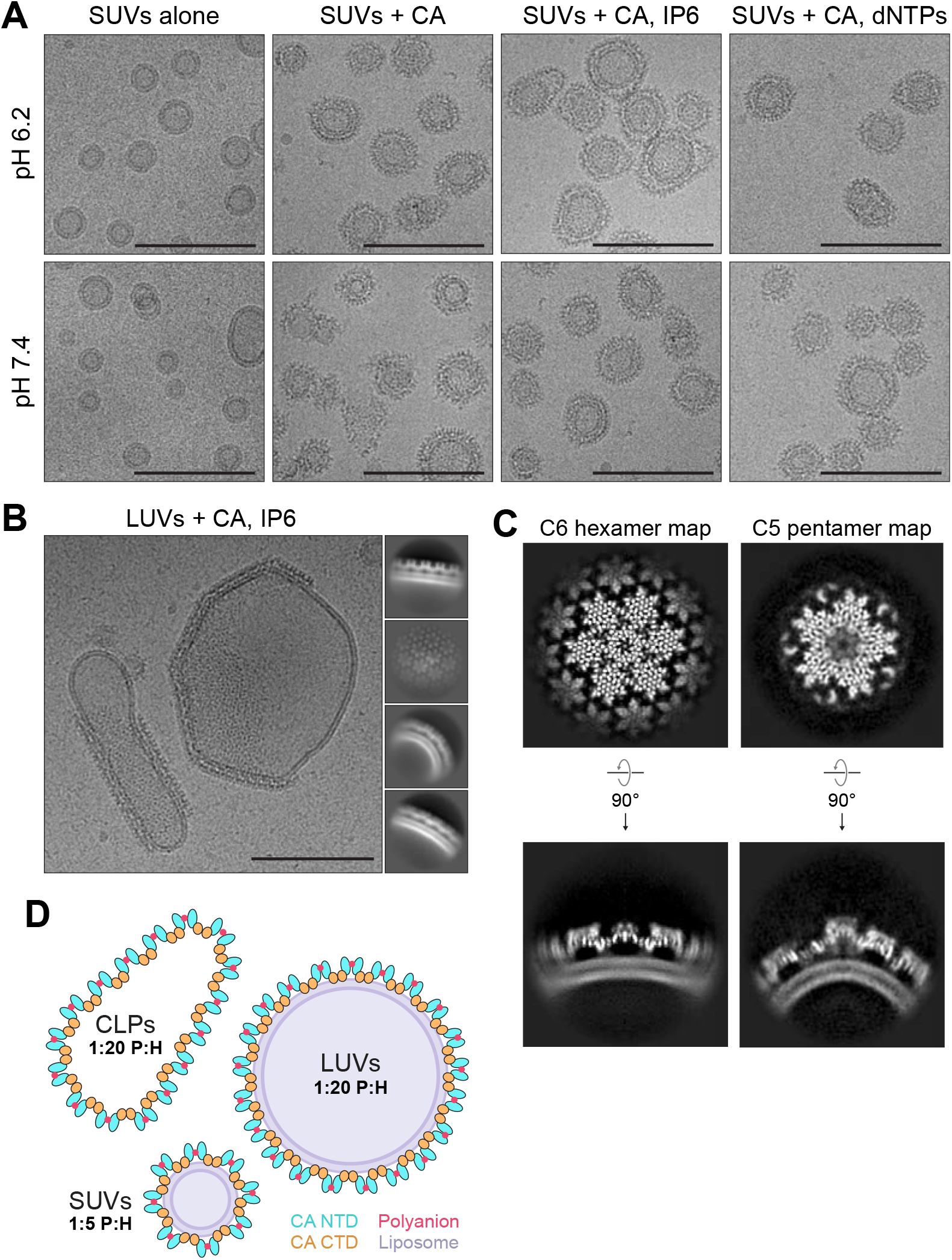
Templating HIV-1 CA lattice on SUVs enriches for pentamer formation when polyanions are present, and the pentamer imposes strict local curvature on the lattice. **A**. Cryo-EM micrographs of small unilamellar vesicles (SUVs) alone or with bound CA-6xHis in the absence or presence of polyanions at pH 6.2 or 7.4. Scale bars, 100 nm. Cryo-EM micrograph of CA-6xHis templated on large unilamellar vesicles (LUVs) at pH 6.2 in the presence of IP6. Scale bar, 100 nm. **C**. Top (top) and side (bottom) orthoslice views of hexamer and pentamer maps from LUV-templated CA-6xHis lattice prepared at pH 6.2 in the presence of IP6. **D**. Comparison of typical observed pentamer-to-hexamer ratios in capsid-like particle (CLP), LUV-templated, and SUV-templated CA lattice assemblies. P, pentamer; H, hexamer.

**SUPPLEMENTAL FIGURE 4.**
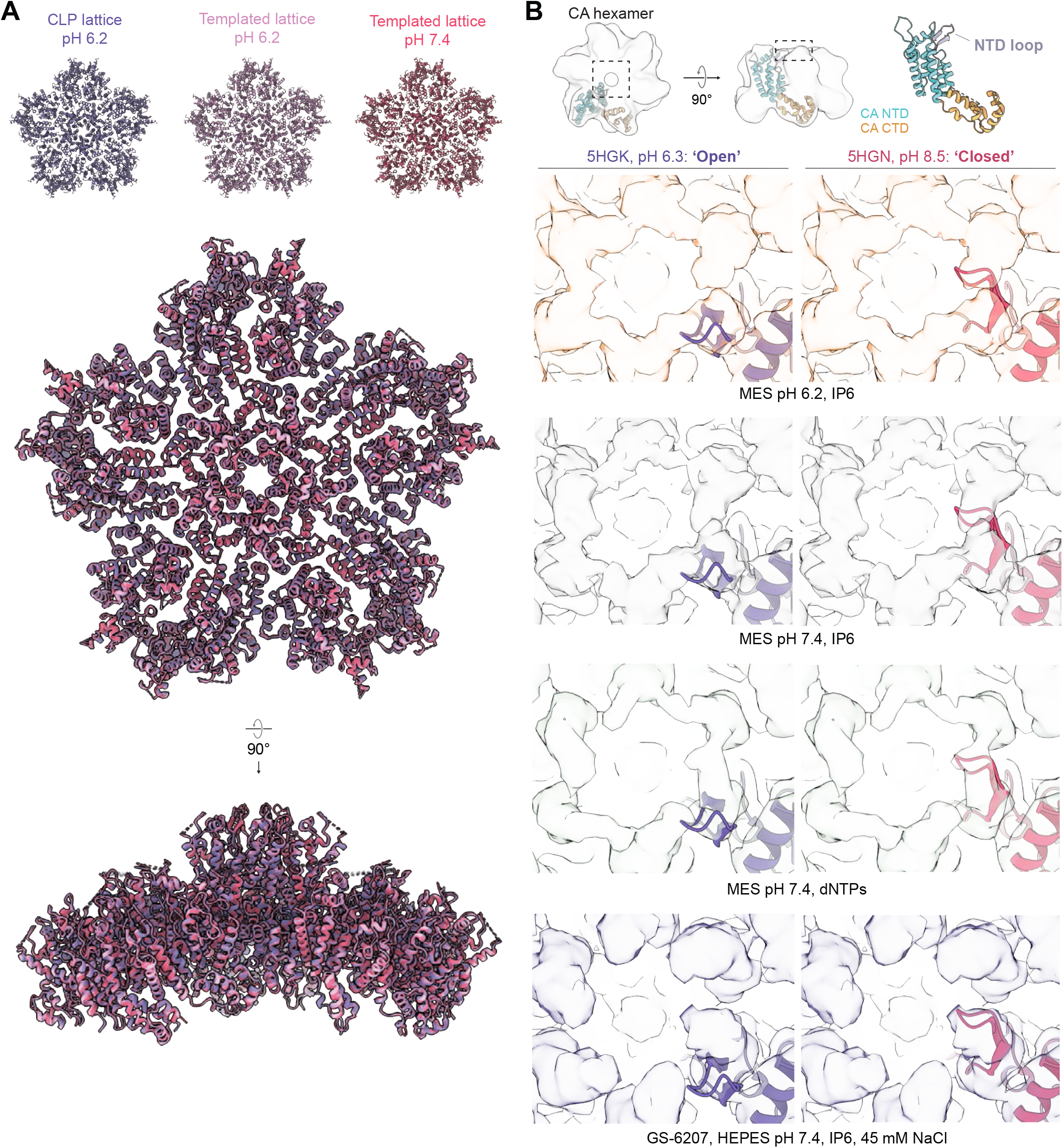
Comparison of CLP and templated CA lattice structures. **A**. Comparison of ‘global’ pentamer atomic models for capsid-like particle (CLP) lattice (purple) and SUV-templated lattice prepared at pH 6.2 (lavender) or pH 7.4 (magenta) structures shown in Figure 2F. **B**. Cryo-EM maps of hexamer central pores with a CA monomer from published crystal structures (Jacques et al., 2016) 5HGK (crystallized at pH 6.3) and 3H4E (crystallized at pH 8.5) docked according to best cross-correlation with the cryo-EM density.

**SUPPLEMENTAL FIGURE 4.**
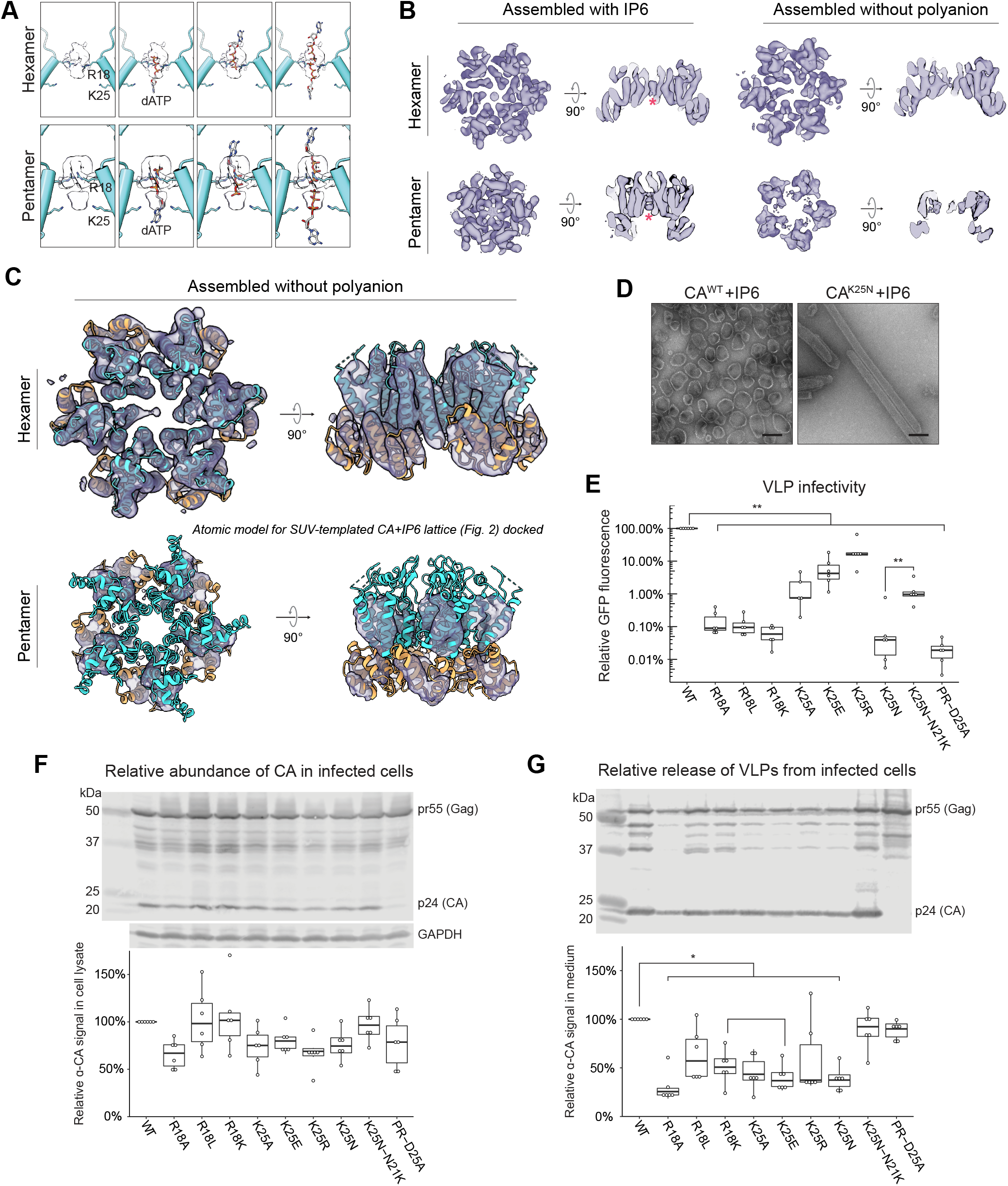
Polyanion binding is critical for proper HIV-1 CA pentamer formation and for virus-like particle infectivity and release from infected cells. **A**. Hypothesized dATP orientations in CA pentamer (top) and hexamer (bottom) pores from structures shown in Figure 3B,D. dATP molecules shown are manually docked into the cryo-EM density to illustrate hypothesized orientations in the pore. **B**. Cryo-EM maps of hexamer and pentamer alone from ‘global’ pentamer maps shown in Figures 3E (left) and 3F (right). Magenta asterisk marks IP6 density. Side views show central pore cross-sections. **C**. Model for templated HIV-1 CA-6xHis lattice prepared at pH 7.4 in the presence of IP6 (presented in Figure 2) docked into the ”Assembled without polyanion” cryo-EM density shown B. **D**. Negative stain TEM micrographs of wild-type (WT) and K25N mutant CA assembled *in vitro* at pH 6.2 with IP6. Scale bars, 100 nm. **E**. Relative infectivity of CA mutants, measured by GFP reporter fluorescence in infected cells. Data are normalized to GAPDH from the western blot shown in F. **F**. Top: representative western blots of HIV CA and GAPDH in infected cell lysates. Bottom: quantification of biological replicates. CA band intensity measurements are normalized to GAPDH. **G**. Top: representative western blots of HIV CA and GAPDH in virus-like particles released from infected cells. Bottom: quantification of biological replicates. CA band intensity measurements are normalized to GAPDH (shown in F). Statistics: N = 6, pairwise comparisons using Wilcoxon rank sum test, * p < 0.05, ** p < 0.01. VLPs, virus-like particles.

