## Supplemental Table 1 for "Structural insights into HIV-1 polyanion-dependent capsid lattice formation revealed by single particle cryo-EM"

| | | Cryo-ET | | SPA | | | | | |
|---|---|---|---|---|---|---|---|---|---|
| | | CLP hex | CLP pent | CLP pent | Templated pent | Templated pent | Templated pent | Templated pent | Templated pent |
| Sample details | Sample pH | 6.2 | 6.2 | 6.2 | 6.2 | 7.4 | 7.4 | 7.4 | 7.4 |
| | IP6 | + | + | + | + | + | | + | |
| | dNTPs | | | | | | + | | |
| | GS-6207 | | | | | | | + | |
| Data collection | Microscope | Titan Krios | Titan Krios | Talos Arctica | Talos Arctica | Talos Arctica | Talos Arctica | Talos Arctica | Talos Arctica |
| | Detector | Gatan K2-XP (Gatan) | Gatan K2-XP (Gatan) | K3 (Gatan) | K3 (Gatan) | K3 (Gatan) | K3 (Gatan) | K3 (Gatan) | K3 (Gatan) |
| | Energy filter | BioQuantum (Gatan) | BioQuantum (Gatan) | BioQuantum (Gatan) | BioQuantum (Gatan) | BioQuantum (Gatan) | BioQuantum (Gatan) | BioQuantum (Gatan) | BioQuantum (Gatan) |
| | Energy filter slit width (V) | 20 | 20 | 20 | 20 | 20 | 20 | 20 | 20 |
| | Nominal magnification | 105,000 | 105,000 | 63,000x | 63,000x | 63,000x | 63,000x | 63,000x | 63,000x |
| | Voltage (kV) | 300 | 300 | 200 | 200 | 200 | 200 | 200 | 200 |
| | Total dose (e-/Å^2) | 123 | 123 | 50 | 50 | 50 | 50 | 50 | 50 |
| | Super-resolution mode? | no | no | yes | yes | yes | yes | yes | yes |
| | Acquisition software | SerialEM | SerialEM | SerialEM | SerialEM | SerialEM | SerialEM | SerialEM | SerialEM |
| | Tilt angle range, step | −60°/60°, 3° (dose-symmetric) | −60°/60°, 3° (dose-symmetric) | - | - | - | - | - | - |
| | Defocus range (μm) | -1.5 to -4.5 | -1.5 to -4.5 | -0.6 to -1.6 | -0.6 to -1.6 | -0.6 to -1.6 | -0.6 to -1.6 | -0.6 to -1.6 | -0.6 to -1.6 |
| | Pixel size (Å) | 1.379 | 1.379 | 1.31 | 1.31 | 1.31 | 1.31 | 1.31 | 1.31 |
| | Frames per tilt or movie | 10 | 10 | 50 | 50 | 50 | 50 | 50 | 50 |
| | Number of tomograms or movies | 66 | 66 | 1,487 | 3,284 | 1,799 | 2,646 | 3,037 | 696 |
| Processing | Final number of particles | 539,700 | 26,220 | 81,428 | 102,495 | 84,603 | 58,577 | 267,942 | 11,558 |
| | Symmetry imposed | C6 | C5 | C5 | C5 | C5 | C5 | C5 | C5 |
| | B factor used for map sharpening (Å) | -230 | -400 | -98 | -100 | -80 | -90 | -103 | -362 |
| | Map resolution at 0.143 FSC (Å) | 3.9 | 6.2 | 3.6 | 3.3 | 3.1 | 3.5 | 3.1 | 7.0 |
| | EMDB ID | | | | | | | | |
| Atomic model | Protein residues | | | 1,379 | 1,379 | 1,379 | 1,379 | 1,379 | - |
| | MolProbity score | | | 1.13 | 1.1 | 0.92 | 1.29 | 1.02 | - |
| | Clash score | | | 3.35 | 3.07 | 1.68 | 4.33 | 2.4 | - |
| | Rotamer outliers (%) | | | 0.26 | 0.17 | 0.43 | 0.26 | 0.14 | - |
| | Ramachandran favored (%) | | | 98 | 98.3 | 98.15 | 97.63 | 1.67 | - |
| | Ramachandran allowed (%) | | | 2 | 1.7 | 1.85 | 2.37 | 98.33 | - |
| | Ramachandran outliers (%) | | | 0 | 0 | 0 | 0 | 0 | - |
| | Ramachandran Z-score | | | 1.16 | 1.62 | 0.52 | 1.47 | 1.3 | - |
| | RMSD, bond length (Å) | | | 0.004 | 0.004 | 0.005 | 0.004 | 0.009 | - |
| | RMSD, bond angles | | | 0.795 | 0.743 | 0.745 | 0.789 | 1.175 | - |
| | PDB ID | | | | | | | | - |
